# *Duox* and *Jak/Stat* signalling influence disease tolerance in Drosophila during *Pseudomonas entomophila* infection

**DOI:** 10.1101/2021.09.23.461578

**Authors:** Arun Prakash, Katy M. Monteith, Mickael Bonnet, Pedro F. Vale

## Abstract

Disease tolerance describes an infected host’s ability to maintain health independently of the ability to clear microbe loads. The Jak/Stat pathway plays a pivotal role in humoral innate immunity by detecting tissue damage and triggering cellular renewal, making it a candidate tolerance mechanism. Here, we find that in *Drosophila melanogaster* infected with *Pseudomonas entomophila* disrupting *ROS*-producing *dual oxidase (duox*) or the negative regulator of Jak/Stat *Socs36E*, render male flies less tolerant. Another negative regulator of Jak/Stat, *G9a* - which has previously been associated with variable tolerance of viral infections – did not affect the rate of mortality with increasing microbe loads compared to flies with functional *G9a*, suggesting it does not affect tolerance of bacterial infection as in viral infection. Our findings highlight that ROS production and Jak/Stat signalling influence the ability of flies to tolerate bacterial infection sex-specifically and may therefore contribute to sexually dimorphic infection outcomes in *Drosophila*.

## 1. Introduction

When organisms experience infection, they face two major challenges to return to a healthy state. The first challenge is to identify and clear the source of the infection. Individuals capable of dealing with the first challenge exhibit low microbe loads because their immune clearance mechanisms are very effective, and are typically labelled ‘resistant’(Boon et al., 2009; Ganz and Ebert, 2010; Lazzaro et al., 2006; Wang et al., 2017). The mechanisms underlying host resistance have been well characterized empirically and often involve the detection of pathogen-derived molecular patterns such as peptidoglycans, and triggering signalling cascades including the immune deficiency (IMD) and Toll pathways, resulting in the downstream expression of antimicrobial peptides (AMPs) that directly kill pathogens (Kleino and Silverman, 2014; Myllymäki et al., 2014; Myllymäki and Rämet, 2014; Palmer et al., 2018; Valanne et al., 2011).

While crucial, pathogen clearance alone will not result in a healthy host, because after pathogen elimination what is left is the tissue damage caused by pathogen growth and as a side-effect of immunopathology. The second challenge to return to healthy state is therefore to repair and regenerate damaged tissues (Martins et al., 2019; Medzhitov et al., 2012; Prakash et al., 2022; Schneider and Ayres, 2008; Soares et al., 2017, 2014). Effective mechanisms of damage signalling and repair may explain why some individuals are tolerant of infection, and are able to experience relatively high health even if their pathogen loads remain high or are not completely cleared (Martins et al., 2019; Soares et al., 2014).

Compared to well-described pathogen clearance mechanisms, we are only beginning to unravel the mechanistic basis of disease tolerance (Martins et al., 2019; Medzhitov et al., 2012; Prakash et al., 2022; Soares et al., 2017, 2014). Likely candidate mechanisms underlying effective tolerance of infection include those that regulate inflammation to reduce immunopathology (Adelman et al., 2013; Cornet et al., 2014; Prakash et al., 2021; Sears et al., 2011); detoxification of host or pathogen derived metabolites (Ferreira et al., 2011; Soares et al., 2017; Vale et al., 2014); or tissue protection and regeneration (Jamieson et al., 2013; Prakash et al., 2022; Soares et al., 2017, 2014). However, the few disease tolerance candidate genes arising from genome-wide association or transcriptomic studies - such as *ghd* (*grainyhead*), *dsb (debris buster), crebA (cyclic response element binding protein) and, dfoxo* (*forkhead box, sub-group O*) - do not appear to be directly associated with classical immune pathways (Dionne et al., 2006; Howick and Lazzaro, 2014; Lissner and Schneider, 2018; Troha et al., 2018).

Here we take advantage of the detailed knowledge of *Drosophila* immunity to investigate the role of damage signalling plays in disease tolerance during systemic bacterial infection. In response to mechanical injury, oxidative stress, and infection, the Jak/Stat pathway is activated by cytokine-like ligands of the unpaired family namely *upd-1, upd-2* and *upd-3* (Agaisse et al., 2003; Chakrabarti et al., 2016; Dostert et al., 2005; Ekengren et al., 2001; Ekengren and Hultmark, 2001; Gilbert et al., 2005; Harrison et al., 1998). *Upd-3* is produced during damage caused by reactive oxygen species (*ROS*), which in turn are produced by *dual oxidase* (*duox*) (Babior, 1995; Klebanoff, 1974; Lee and Kim, 2014). The extracellular binding of *upd-3* to *Domeless* (*dome*), leads to the phosphorylation of *Hopscotch* (*hop*). This then leads to the phosphorylation of *Stat92E*, and its translocation to the nucleus (Myllymäki and Rämet, 2014). In the nucleus, in addition to the production of factors that are necessary for repairing cellar damage, *Stat92E* also induces the expression of *Socs36E*, a negative regulator of *Hopscotch* (Kiu and Nicholson, 2012). Recent work has also highlighted the role of the histone H3 lysine 9 methyltransferase (also called *G9a*) in negatively regulating the expression of the Jak/Stat pathway during infection (Merkling et al., 2015).

Focusing on its role in immunity, there is substantial evidence that Jak/Stat signalling plays a key role in wound healing, gut immunity, and downstream AMP production (Chakrabarti et al., 2016; Kemp et al., 2013; Lamiable and Imler, 2014; Tafesh-Edwards and Eleftherianos, 2020). For instance, during enteric bacterial infection in *Drosophila*, the Jak/Stat pathway contributes to intestinal immunity by regulating intestinal stem cell (ISC) proliferation and epithelial cell renewal via epidermal growth factor (*EGFR)* signalling (Buchon et al., 2010; Chakrabarti et al., 2016; Ohlstein and Spradling, 2006). The absence of epithelial renewal leads to a loss of structural integrity and increased susceptibility to bacterial infections (Buchon et al., 2009). In cellular immunity, Jak/Stat signalling is central to the production, differentiation and maintenance of blood cells in insects (Banerjee et al., 2019; Meister and Lagueux, 2003). The Jak/Stat pathway is also important in humoral immunity to viral infection (Dostert et al., 2005), where a loss of regulation of Jak/Stat by the epigenetic negative regulators *G9a* results in reduced tolerance of *Drosophila C virus* infections due to increased immunopathology (Merkling et al., 2015). This specific result motivated us to question whether the effects of *G9a*-mediated Jak/Stat regulation on tolerance were specific to viral infection, or if the regulation of Jak/Stat also affects disease tolerance during bacterial infection.

We investigated the tolerance response of *Drosophila* during septic infection with the bacterial pathogen *P. entomophila*, using transgenic flies lacking various components of Jak/Stat signalling and regulation. Further motivated by the widespread observation of sexually dimorphic immunity *reviewed in* (Belmonte et al., 2020; Klein and Flanagan, 2016) and particularly that the effects of *G9a* on tolerance of *DCV* infection are more pronounced in female flies (Gupta and Vale, 2017; Merkling et al., 2015), we also focused on assessing sex differences in how Jak/Stat signalling affects tolerance of *P*.*entomophila* infection.

## 2. Materials and methods

### 2.1 Fly strains and maintenance

We used several *D*. melanogaster transgenic lines with TE mobilization using a P-element construct and subsequent loss-of-function for *Duox -* P{SUPor-P}Duox^KG07745^ (Hurd et al., 2015), *Domeless -* P{SUPor-P}^KG08434^, *Hopscotch* - P{SUPor-P}hop^KG01990^(Bellen et al., 2004), *Socs36E -* P{EPgy2}Socs36EEY06665 (Monahan and Starz-Gaiano, 2013). All lines were on the *yw* background (Eleftherianos et al., 2014) which served as a control genotype (detailed information is presented in **Fig** S1 and S2 and Table S1). We also used *G9a* mutant flies (that is, *G9a*^-/-^, also known as *G9a*^*DD2*^ generated previously by mobilization of the P-element *KG01242* located in the 5’ UTR of the gene(Kramer et al., 2011)) and control *G9a*^*+/+*^ (Merkling et al., 2015). We maintained all the fly lines in a 12ml plastic vials on a standard cornmeal diet *see* (Siva-Jothy et al., 2018), at 25°C (±2°C). We used 3-5-day-old adult flies for all our experiments (see below). First, we housed 2 males and 5 females for egg laying (48 hours) in a vial containing fresh food. We then removed the adults and the vials containing the eggs were kept in 25°C incubator for 14 days, or until pupation. We placed the newly eclosed individuals (males and females separately) in fresh food vials until the experimental day (3 days).

### 2.2 Bacterial culture preparation

We used *P. entomophila* cultured overnight in Luria broth (LB) at 37°C under constant agitation that is, 120 revolutions per minute (rpm). *P. entomophila* is a gram-negative bacterium naturally found in soil and aquatic environments, known to be highly pathogenetic for *D. melanogaster* (Dieppois et al., 2015; Vodovar et al., 2005). Upon reaching 0.75 OD_600_ we pelleted the culture by centrifuging during 5 minutes at 5000rpm at 4°C, and then removed the supernatant. We resuspended the bacteria in 1xPBS (phosphate buffer saline) and prepared the final infection inoculum of OD_600_ of 0.05 for all our infection assays.

### 2.3 Systemic infection assay

We used a split-vial experimental design (see ***Fig***. *S3*), where, after infection, each vial containing 25 flies (of each sex and fly line combination) were divided into 2 vials for measuring survival following infection (n= 15 vials of 15-17 flies/vial/infection treatment/sex/fly line) and internal bacterial load (n= 15 vials of 8-10 flies/vial/infection treatment/sex/fly line). With this split-vial design we were able to use replicate-matched data for both survival and bacterial load to estimate disease tolerance for each fly line (that is, for each replicate group, mean fly survival with respect to mean internal bacterial load). We infected 3-5-day old male and female adult flies using a 0.14mm insect minutein needles bent at 90° angle to avoid damaging the internal tissues by dipping in *P. entomophila* bacterial inoculum of OD_600_ of 0.05, resulting in 50-70 bacterial cells/fly. For mock controls we substituted bacterial solution with sterile 1xPBS. After stabbed the flies in the sternopleural region of the thorax (Khalil et al., 2015). We then placed males and females separately onto fresh food vials and incubated at 25°C. We scored the flies (both infected and control) every 2-3 hours for the first 48-hours following infection, then 2-3 times each day for the next 6 days (150 hours).

### 2.4 Measuring bacterial load

To quantify internal bacterial load after 24-hours following systemic *P. entomophila* infection first, we thoroughly washed each fly with 70% ethanol for 30 sec to surface sterilize and then rinsed twice with autoclaved distilled water. We plated the second wash on LB agar plates and confirmed that the surface bacteria were successfully removed after sterilization. We then transferred individual fly onto 1.5ml micro centrifuge tubes and homogenized using a motorized pestle for approximately 30-60 seconds in 100µl LB broth (n=30 fly homogenates/sex/infection treatment/ fly line). We performed serial dilution of each fly homogenate up to 10^−6^ fold and added 4μL aliquot on a LB agar plate. We incubated the plate overnight for 18h at 30°C and counted the resultant bacterial colonies manually (Siva-Jothy et al., 2018). We note that mock-infected control fly homogenates did not produce any colonies on LB agar plates.

### 2.5 Statistical analyses

#### 2.5.1 Survival

We analysed the survival data with a Cox mixed effects model using the R package ‘coxme’ (*Therneau 2015*) for different treatment groups (*P. entomophila* systemic infection and mock controls) across males and females. We specified the model as: survival ∼ fly line * treatment * sex * (1|vials/block), with ‘fly line’, ‘treatment’ and ‘sex’ and their interactions as fixed effects, and ‘vials’ nested within a ‘block’ as a random effect.

#### 2.5.2 Bacterial load

We found that the bacterial load data were not normally distributed (tested with Shapiro–Wilks’s test for normality). We therefore used a non-parametric one-way ANOVA Kruskal-Wallis test to test the effects of each fly line and sex on internal bacterial load.

#### 2.5.3 Measuring disease tolerance

We analysed disease tolerance as the linear relationship between fly survival against bacterial load (Ayres and Schneider, 2012; Louie et al., 2016; Oliveira et al., 2020; Raberg et al., 2007). To this end, we employed ANCOVA by fitting ‘fly line’ and ‘sex’ as categorical fixed effects and ‘bacterial load’ as a continuous covariate, and their interactions as fixed effects. Since we were interested in identifying how each transgenic line differed from the control line, we compared the estimates of the model slope using pairwise comparison (f-test; *yw* vs. different transgenic lines) to test the extent to which each transgenic line significantly differed from the control in tolerating bacterial infections.

## 3. Results and Discussion

### 3.1. Following systemic bacterial infection, disruption of *Duox* or different components of Jak/Stat pathway result in variable survival outcomes

Overall, we found that disruption of *Duox* or the Jak/Stat pathway (either by disrupting the positive regulators *upd3* and *domeless*, or overactivation by disrupting the negative regulator *socs36E*) affected fly survival during bacterial *P. entomophila* infections (***Fig***. *1A* and *B*, ***Table*** *1* and *SI-2*). Both male and female flies lacking *duox* (*ROS* producing *dual oxidase*) were more susceptible to *P. entomophila* infections compared to the control line (*yw*) (***Fig***. *1A* and *B*, ***Table*** *1* and *SI-2*). However, other transgenic lines showed slightly improved survival relative to the functional control line. These included male and female flies lacking the transmembrane receptor *domeless*, and males lacking the negative regulator *Soc36E* (see hazard ratio in ***Fig***. *1B*, ***Table*** *1* and *SI-2*).

**Figure 1.**
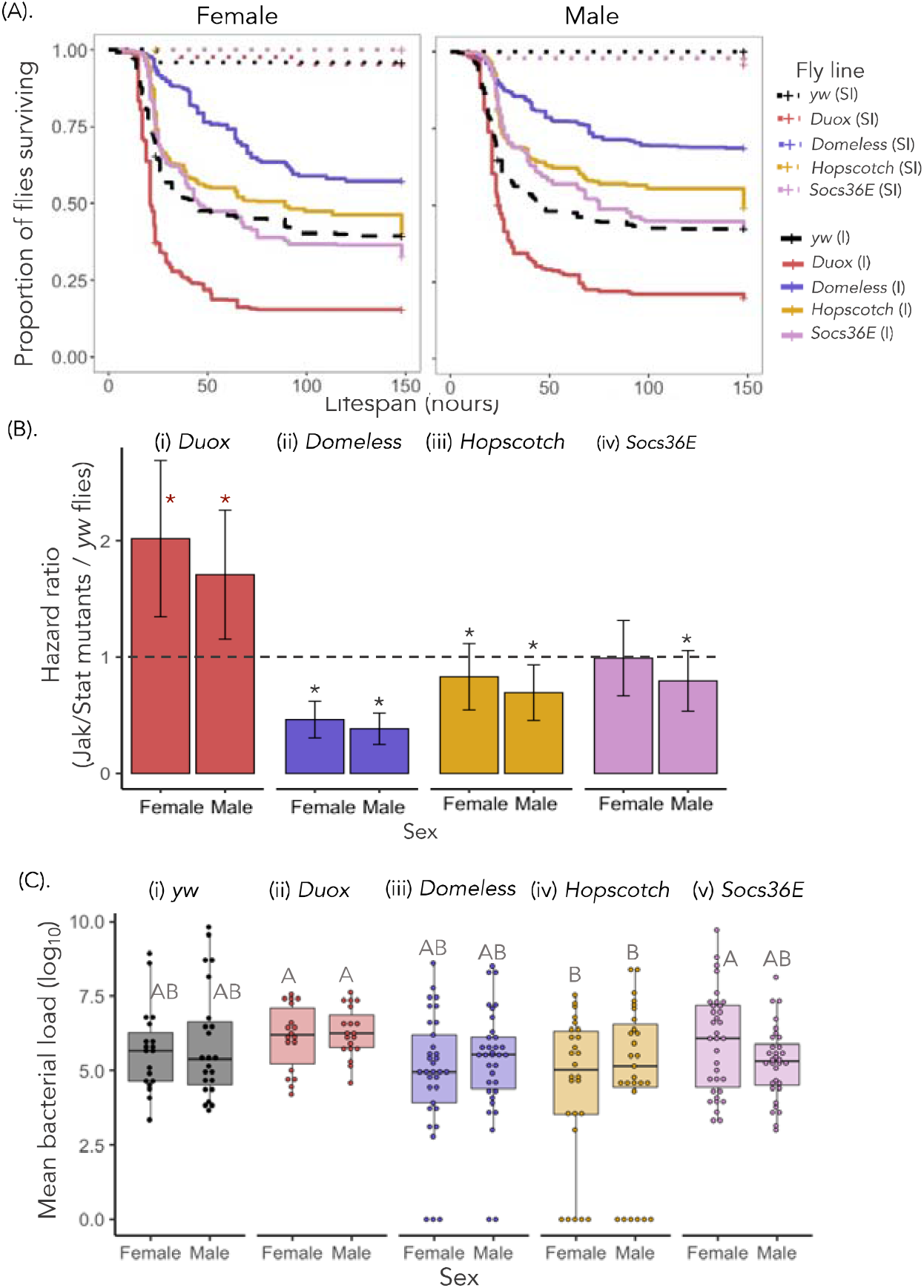
**(A)** Survival curves for control *yw* flies and flies lacking Jak/Stat pathway components for females and males exposed to systemic *P. entomophila* of infection dose OD_600_=0.05 (n= 15 vials with 15-17 flies each vial/fly line/treatment/sex/infection dose). [‘*’ indicates that the Jak/Stat transgenic lines are significantly different from *yw* flies]. **(B)** Estimated hazard ratios calculated from the survival curves for males and female flies (y*w* and with flies lacking components of Jak/Stat signalling and *duox*). A greater hazard ratio (>1) indicates higher susceptibility of Jak/Stat mutants than control while (<1) indicates transgenic lines have better survival than control flies to systemic bacterial infection (p=<0.05). **(C)** Bacterial load (mean log_10_) measured 24 hours following infection (n= 15 vials with 8-10 flies each vial/fly line/treatment/sex/infection dose). [significantly different fly lines are connected by different letters using Tukey’s HSD as a post hoc analysis of pairwise comparisons].

**Table 1:** Summary of estimated hazard ratio from the cox proportional model. A greater hazard ratio estimates (>1) indicates that Jak/Stat mutant flies are more susceptible to *P. entomophila* infection than *yw* control flies while lower ratio (<1) indicates that transgenic lines have better survival than *yw* control.

| *Fly line* | *sex* | *estimate* | *P* | *lower 95%* | *upper 95%* |
| --- | --- | --- | --- | --- | --- |
| *Domeless* | Female | 0.462 | <0.001 | 0.391 | 0.548 |
|  | Male | 0.383 | <0.001 | 0.322 | 0.457 |
| *Duox* | Female | 2.017 | <0.001 | 1.712 | 2.384 |
|  | Male | 1.707 | <0.001 | 1.455 | 2.009 |
| *Hopscotch* | Female | 0.830 | 0.03 | 0.701 | 0.986 |
|  | Male | 0.694 | <0.001 | 0.585 | 0.824 |
| *Socs36e* | Female | 0.990 | 0.91 | 0.843 | 1.167 |
|  | Male | 0.795 | 0.006 | 0.676 | 0.937 |

### 3.2 Control *yw* and Duox / Jak/Stat transgenic deletion lines exhibit similar bacterial loads

We investigated whether the variation we observed between transgenic lines in mortality could be explained by differences in their bacterial load. Given that most mortality occurred just after 24 hours for most of our fly genotypes (***Fig***. *1A*) we quantified bacterial load at 24 hours following infection. Both control and transgenic lines exhibited similar levels of bacterial load 24 hours following infection with *P. entomophila* (***Fig***. *1C*, ***Table*** *SI-3*). Therefore, despite no substantial difference in microbe loads at 24-hours post infection, transgenic lines showed variable survival. This would fit the functional definition of disease tolerance as for the same bacterial load some lines appear to be more tolerant (survive longer, such as *domeless*) while others are less tolerant (e.g., *duox*).

### 3.3 Disrupted expression of *Duox* or *Jak-Stat* signalling leads to differences in disease tolerance phenotypes

While the results above are indicative of variable tolerance depending on the Jak/Stat disruption, we carried out a formal analysis of disease tolerance using the slope of the linear reaction norm between fly survival and microbe load, where each data point is the matched survival / CFU data for one replicate vial (see methods and ***Fig***. *S3* for description of split-vial design). Here, the differences in tolerance between Jak/Stat deletion and the control fly line are indicated by a significant interaction between the bacterial load and the fly line for survival, which reflects the overall rate at which fly health (survival) changes with bacterial load between fly lines. Overall, we found that the transgenic lines showed differences in disease tolerance phenotypes compared to control in both males and females, and this effect was driven mainly the Duox-deficient lines, which showed a much steeper decline in survival with increasing *P. entomophila* bacterial loads (***Fig***. *2A* *and 2B*, ***Tables 2 and*** *3*). Given the role of *duox* in producing *ROS*, one possible explanation for decreased tolerance in the *duox* transgenic line is flies require intracellular *ROS* (oxidative burst) such as *H*_*2*_*O*_*2*_ (hydrogen peroxide) for the activation of cellular reponses during wounding and injury, in addition to Toll and Jak/Stat activation (Chakrabarti and Visweswariah, 2020). In other work, wild type (*w*^*Dahomey*^) males showed higher levels of *duox* expression and *ROS* following *Ecc* (*Erwinia catovora*) infection (Regan et al., 2016), which may suggest that loss of function of *duox* might impact males more than females, as observed in this experiment (Fig. 2B).

**Figure 2.**
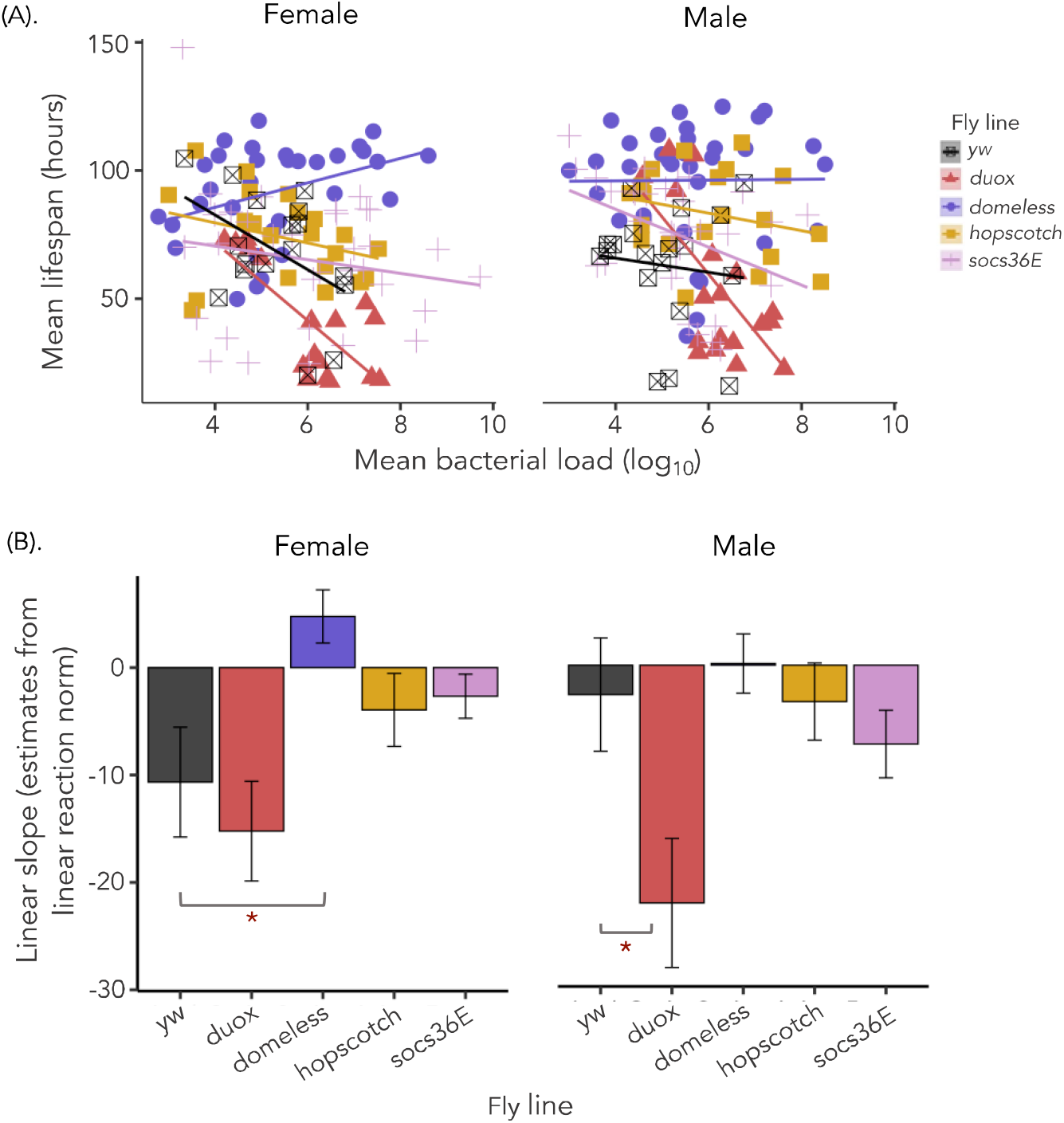
**(A)**. The relationship between fly survival (measured as mean lifespan) and mean bacterial load (as mean CFUs - Colony Forming Units) analysed using linear models for female and male flies. Each point shows data for median lifespan and mean CFUs of 15 vials (with each vial containing 25 flies/sex/fly line combination after 24 hours post systemic bacterial exposure. The data shown here are for the infection doses (OD_600_=0.05). **(B)**. Represents estimates of negative slope of the linear reaction norm extracted from the linear models. [Maroon asterisks ‘*’ on the lower side of the panel B indicates that transgenic lines are significantly different from control *yw*, analysed using the F-test pairwise comparisons of estimates of the linear reaction norm for both males and females separately (see Table-3)]. Grey asterisks ‘*’ on the upper side of the panel B indicates sex differences within the fly line that is, males and females significantly differ in tolerance to systemic bacterial *P. entomophila* infection.

**Table 2.** Summary of ANCOVA. To assess differences in infection tolerance (fly survival with increasing bacterial burden) following systemic *P. entomophila* infection with OD_600_=0.05 infection dose, 24 hours following infection. We analysed ANCOVA and fitted ‘sex’ as categorical fixed effects, ‘mean bacterial load (log_10_)’ as a continuous covariate and their interactions as fixed effects for the transgenic lines.

| *Fly line* | *Source* | *DF* | *Sum of Sq.* | *F ratio* | *P* |
| --- | --- | --- | --- | --- | --- |
| ***Female*** | Fly line | 4 | 24817.1 | 15.27 | **<0.001** |
|  | Bac. load | 1 | 4482.9 | 11.03 | **0.0012** |
|  | Fly line X bac. load | 4 | 7642.8 | 4.7 | **0.0015** |
| ***Male*** | Fly line | 4 | 16964.8 | 9.22 | **<0.001** |
|  | Bac. load | 1 | 6122.6 | 13.32 | **0.0004** |
|  | Fly line X bac. load | 4 | 5737.5 | 3.12 | **0.017** |

**Table 3:** Summary of F-test pairwise comparisons of estimates of linear slopes (from the linear model) transgenic lines compared to the *yw* control.

| *sex* | *line* | *SSE* | *ddf* | *slope diff* | *std err* | *F ratio* | *p* |
| --- | --- | --- | --- | --- | --- | --- | --- |
| **Female** | *Duox* | 10018.68 | 30 | -13.15 | 3.09 | 0.53 | 0.47 |
|  | *Domeless* | 16075.94 | 45 | 1.83 | 2.25 | 8.38 | **0.0058** |
|  | *Hopscotch* | 11897.99 | 34 | -5.99 | 2.64 | 1.39 | 0.24 |
|  | *Socs36E* | 27135.53 | 47 | -3.77 | 2.28 | 1.48 | 0.22 |
| **Male** | *Duox* | 17512.77 | 34 | -11.25 | 4.47 | 5.25 | **0.028** |
|  | *Domeless* | 27106.63 | 49 | -0.46 | 2.67 | 0.19 | 0.65 |
|  | *Hopscotch* | 15019.55 | 36 | -3.2 | 2.8 | 0.01 | 0.91 |
|  | *Socs36E* | 22285.12 | 47 | -6.17 | 2.75 | 0.54 | 0.46 |

An unexpected observation was that flies lacking *domeless* showed slightly increased survival relative to the *yw* control (Fig 1) (and a trend for increased tolerance, though not statistically significant, Fig 2). Given the role of *domeless* as an activator of Jak-Stat signalling, this might suggest that Jak/Stat activation may be costly to flies. While immune deployment and regulation is highly energy demanding across most species (McKean et al., 2008; Nystrand and Dowling, 2020; Schwenke et al., 2016; Vale et al., 2015), the physiological costs of specific individual immune components and pathways remains understudied and an open question for future research.

### 3.4 Disruption of *G9a* does not affect tolerance of *P*.*entomophila*

The negative regulator of Jak/Stat, *G9a*, was previously identified as being important for tolerating *Drosophila C Virus* (DCV) infections (Merkling et al., 2015). Subsequent work exploring sex differences in this response found that *G9a*^*+/+*^ (control) females had higher tolerance than *G9a*^*−/−*^ females, when measured across a range of viral DCV doses (Gupta and Vale, 2017). We wanted to test whether the loss of function of *G9a* also affects fly survival and disease tolerance in response to bacterial infections. Overall, we found that loss of *G9a* makes both males and females more susceptible to *P. entomophila* infections, (***Fig***. *3A* for survival and ***Fig***. *3B* for hazard ratio, ***Table*** *4* and ***Table*** *SI-4*). To test if this increased mortality in *G9a*^*-/-*^ flies was associated with higher bacterial replication we measured bacterial load following 24 hours *P. entomophila* systemic infection. We found that *G9a*^*-/-*^ females exhibited higher bacterial load than *G9a*^*+/+*^ (control) flies, while males showed similar bacterial load as *G9a*^*+/+*^ flies (***Fig***. *3C*, ***Table*** *SI-5*). However, the overall ability to tolerate *P. entomophila* bacterial infections (that is, measured as *G9a* fly’s survival relative to its bacterial load) remained similar across both males and females *G9a* flies that is, both *G9a*^*-/-*^ and *G9a*^*+/+*^ controls (***Fig***. *3D*, ***Table*** *5*, and ***Table*** *6* for comparison between estimates of tolerance slope). Thus, despite the previously identified role of this negative regulator of Jak/Stat in tolerating viral infections by reducing immunopathology (Gupta and Vale, 2017; Merkling et al., 2015), *G9a* does not appear to affect bacterial disease tolerance in either sex.

**Figure 3.**
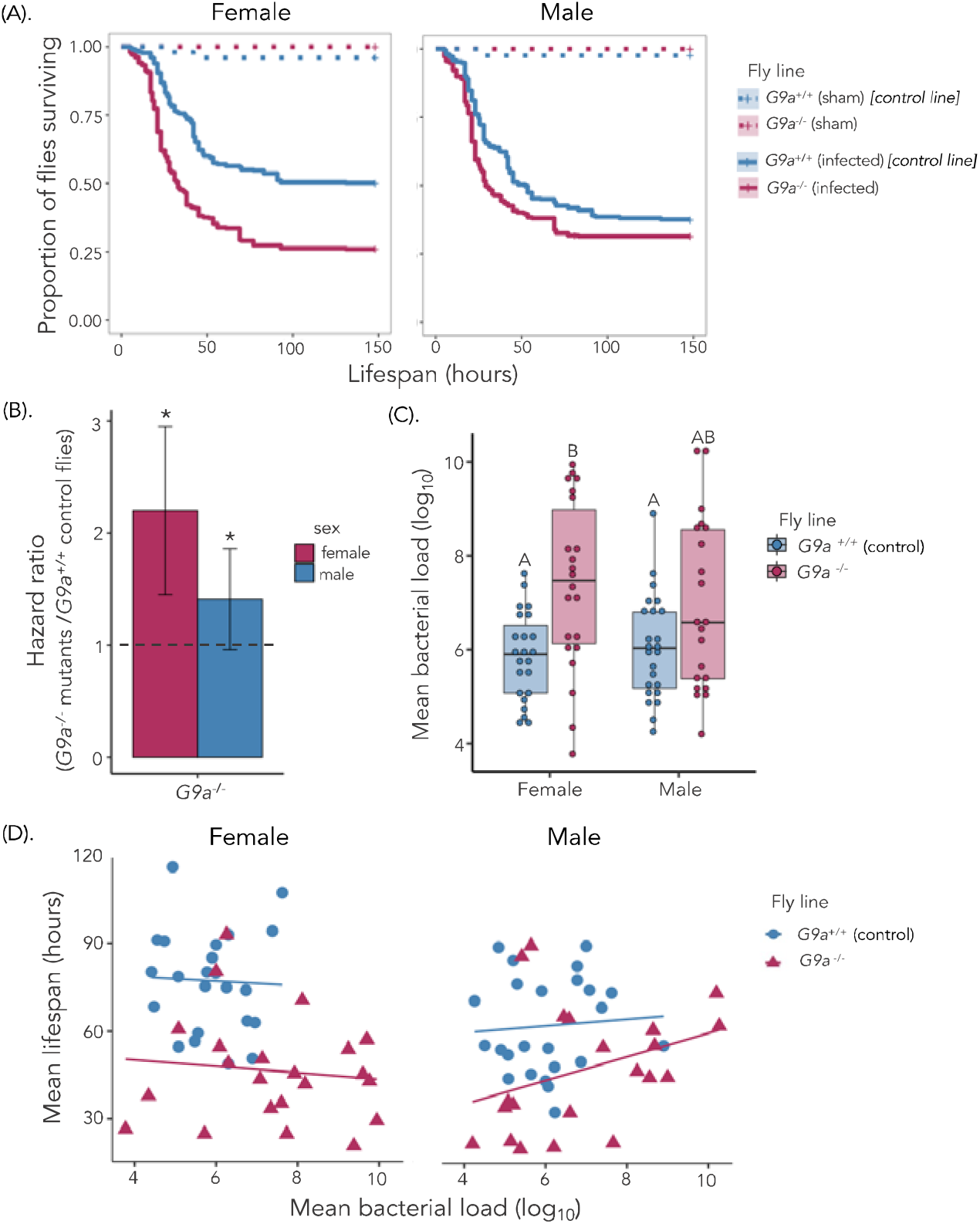
**(A)** Survival curves for control *G9a*^*+/+*^ flies and *G9a*^-/-^ flies lacking *G9a* the epigenetic regulator of Jak/Stat for female and male flies exposed to systemic *P. entomophila* of infection dose OD_600_=0.05 [n=15 vials with 15-17 flies in each vial/fly line/treatment/sex]. **(B)** Estimated hazard ratios calculated from the survival curves for males and female flies (control y*w* and flies without *G9a*). A greater hazard ratio (>1) indicates higher susceptibility of *G9a*^-/-^ to bacterial infection relative to control flies. [‘*’ indicates that the *G9a*^*-/-*^ flies are significantly different from *G9a*^*+/+*^ flies]. **(C)** Bacterial load (mean log_10_) measured 24 hours following infection (n=15 vials with 8-10 flies in each vial/fly line/ treatment and sex combination). [Significantly different fly lines are connected by different letters using Tukey’s HSD as a post hoc analysis of pairwise comparisons]. **(D)** Linear tolerance to *P. entomophila* infection – the relationship between *G9a* fly survival (measured as mean lifespan) and bacterial load (as mean CFUs - Colony Forming Units) analysed using linear models for female and male flies (both *G9a*^*-/-*^ and *G9a*^*+/+*^).

**Table 4:** Summary of estimated hazard ratio from the cox proportional model. A greater hazard ratio (>1) indicates that *G9a*^*-/-*^ flies are more susceptible to *P. entomophila* infection than control (*G9a*^*+/+*^) flies.

| *sex* | *Fly line* | *estimate* | *p* | *Std err* |
| --- | --- | --- | --- | --- |
| Female | G9a-/- | 2.2 | <0.001 | 0.75 |
| Male | G9a-/- | 1.41 | <0.001 | 0.45 |

**Table 5.** Summary of ANCOVA. To assess differences in infection tolerance (fly survival with increasing bacterial burden) following systemic *P. entomophila* infection with OD_600_=0.05 infection dose, 24 hours following infection. We analysed ANCOVA and fitted ‘sex’ as categorical fixed effects, ‘mean bacterial load (log_10_)’ as a continuous covariate and their interactions as fixed effects for each of the fly lines (*G9a*).

|  | *Fly line* | *Source* | *DF* | *Sum of Sq.* | *F ratio* | *p* |
| --- | --- | --- | --- | --- | --- | --- |
| **G9a** | **Female** | Fly line | 1 | 6802.2 | 20.21 | **<0.001** |
|  |  | Bac. load | 1 | 53.19 | 0.158 | 0.69 |
|  |  | Fly line X bac. load | 1 | 1.610 | 0.004 | 0.94 |
|  | **Male** | Fly line | 1 | 3042.5 | 8.685 | **0.005** |
|  |  | Bac. load | 1 | 533.13 | 1.521 | 0.22 |
|  |  | Fly line X bac. load | 1 | 166.21 | 0.474 | 0.49 |

**Table 6:** Summary of F-test pairwise comparisons of estimates of the linear slopes (linear reaction norm) for *G9a* ^*-/-*^*relative to G9a* ^*+/+*^ control fly lines.

| *Sex* | *Fly line* | *Fly line* | *F Ratio* | *p* |
| --- | --- | --- | --- | --- |
| Female | *G9a*<sup>-/-</sup> | *G9a*<sup>+/+</sup> | 0.005 | 0.94 |
| Male | *G9a*<sup>-/-</sup> | *G9a*<sup>+/+</sup> | 0.474 | 0.49 |

## 4. Concluding remarks

Tissue damage signalling and repair mechanisms such as Jak/Stat are important from a therapeutic perspective because they have the potential to boost host tolerance by minimising disease severity (Soares et al., 2014; Vale et al., 2016). Our data show that loss of Jak/Stat pathway components reduces overall survival following *P*.*entomophila* infection and that this is not caused by impaired pathogen clearance but due to lower disease tolerance. These observations have parallels in human infection. For instance, dysregulation of cytokines and interferons (JAK signalling - Tyrosinekinase2) result in immunodeficiency while defective STAT increases the risk of autoimmunity (O’Shea et al., 2014, 2013). Drugs that inhibit JAK have been shown to be effective in treating several autoimmune diseases by targeting cytokine-dependent pathways, while STAT inhibitors have been promising candidates in the context of cancer (Miklossy et al., 2013; Pérez-Jeldres et al., 2019; Salas et al., 2020). It may therefore be possible to repurpose these existing drugs to improve host tolerance of infection. In summary, our work highlights that Jak/Stat directly impacts the ability to tolerate bacterial infection and that this response differs between males and females. Jak/Stat mediated disease tolerance may be a potential source of sexually dimorphic response to infection in *Drosophila*.

## 5. Acknowledgments

We thank Ashworth fly group members for helpful discussion, and Angela Reid, Lucinda Rowe, James King and Alison Fulton for help with media preparation. We acknowledge funding and support from the Branco Weiss fellowship and a Chancellor’s Fellowship to PFV; a Darwin Trust PhD studentship to AP from the School of Biological Sciences, The University of Edinburgh. For the purpose of open access, the author has applied a Creative Commons Attribution (CC BY) licence to any Author Accepted Manuscript version arising from this submission.

## Notes

https://zenodo.org/record/7867071#.ZEk7yezMLPY

### Competing Interest Statement

The authors have declared no competing interest.

### Summary of Updates

- qPCR validation of the transgenic lines is now provided in Supplementary material - Some re-writing of the manuscript to improve clarity and correct various typographical errors in the previous version. - New title to more accurately reflect the main findings

