## Supplementary Figures and Tables for "*Duox* and *Jak/Stat* signalling influence disease tolerance in Drosophila during *Pseudomonas entomophila* infection"

Figure S1. Comparison of yw to an outcrossed population

Figure S2. qPCR confirmation of gene disruption in transgenic flies.

Figure S3: Experimental design.

Tables S1 – S6. -Summaries of statistical analyses.

***Figure S1. Comparison of yw to an outcrossed population* (A)** Survival curves for males and females of *yw* or an advanced outcross population exposed to systemic *P. entomophila* of infection dose OD_600_=0.05 (n=20-30 flies/vial (15 vials)/fly line/treatment/sex/infection dose). **(B)** Estimated hazard ratios calculated from the survival curves for males and female flies (*Outcross* and *yw* flies). A greater hazard ratio (>1) indicates higher susceptibility to bacterial infection. [‘n.s’ in the panel indicates that *Outcross* flies are not significantly different from *yw* flies]. **(C)** Internal bacterial load (as mean CFUs- colony forming units) measured around 24 hours post systemic *P. entomophila* infection. [significantly different fly lines are connected with different letters using Tukey’s HSD as a post hoc analysis of pairwise comparisons].

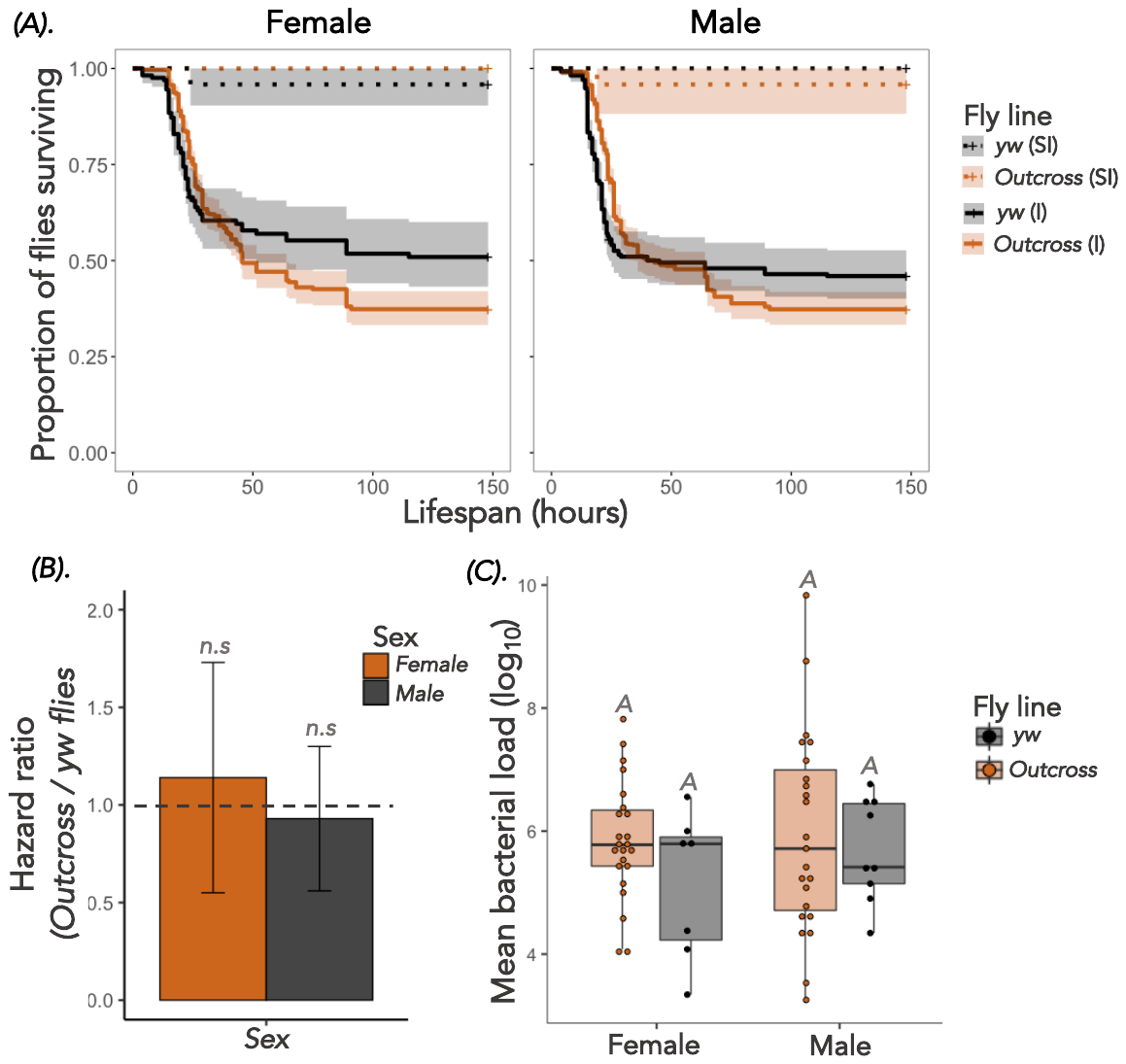

**Both wildtype (*yw*) and *Outcrossed* flies show similar response to systemic *P. entomophila* infection:** All our transgenic lines were on a *yw* genetic background. Previous work has shown that *yw* flies have lower basal levels of nitric oxide and are more susceptible to infection compared to other lab wild types such as *w^1118^, Oregon R, Canton-S (Eleftherianos et al., 2014)*. Before addressing whether the *Jak/Stat* pathway is involved in disease tolerance of bacterial infection, we wanted to better understand how wild type (*yw*) flies survive *P. entomophila* infections relative to other wild-type strains. We used an *Outcrossed* fly population as another control for comparison, which was originally created using 100 pairwise crosses of 113 *DRGP* lines and maintained as an outcrossed population for over 19 generations [see (Savola et al., 2021). We found that males and females of both wild type *yw* and the *Outcross*ed flies showed comparable survival following *P. entomophila* systemic infection and exhibited similar levels of internal bacterial loads when measured 24-hours post-infection. Since both the wildtype (*yw*) and *Outcross* fly lines showed similar infection responses to *P. entomophila*, all remaining experiments only used *yw* as the wildtype control line.

***Figure S2. qPCR confirmation of gene disruption in transgenic flies.*** To confirm the status of each fly line with disrupted Duos or Jak-Stat gene, we measured the expression of TotA and Upd3, two Jak-Stat-responsive genes. Upd3 is also usually upregulated because of ROS-mediated damage. ANOVA for gene expression is reported below as are qPCR conditions for each gene of interest GOI).

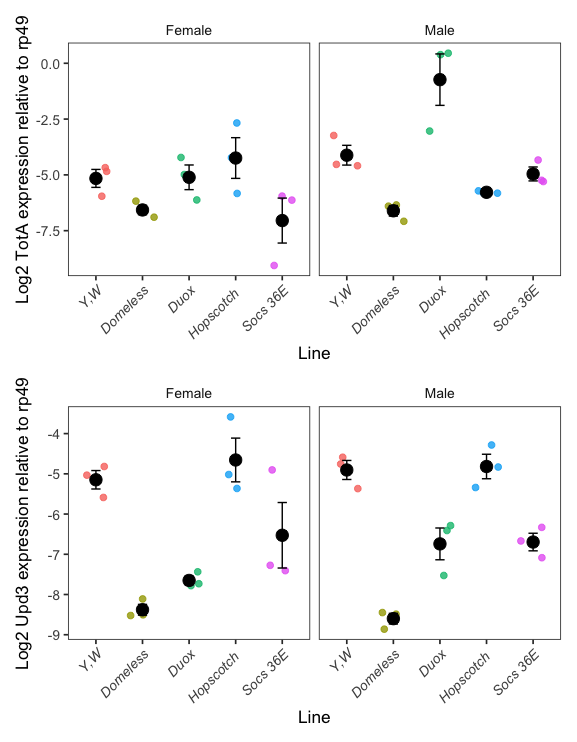

***Table S1 ANOVA summary***

| **TotA** | Df | Sum Sq | Mean Sq | F value | Pr(>F) |
| --- | --- | --- | --- | --- | --- |
| Fly line | 4 | 47.88 | 11.97 | 9.89 | 1.39E-04 |
| Sex | 1 | 10.55 | 10.55 | 8.72 | 0.008 |
| Sample:Sex | 4 | 29.87 | 7.47 | 6.17 | 0.002 |
| **UPD3** | Df | Sum Sq | Mean Sq | F value | Pr(>F) |
| Fly line | 4 | 58.20 | 14.55 | 34.37 | 1.06E-08 |
| Sex | 1 | 0.11 | 0.11 | 0.26 | 0.619 |
| Sample x Sex | 4 | 1.37 | 0.34 | 0.81 | 0.533 |

***qPCR conditions***

Each plate had 4 x non-RT samples and 2x blank controls for GOI primer

Several plates were run to ensure primer optimisation (7 in total) but all data was collected from 4 separate plates (2 plates for TotA and 2 plates for UPD3)

**TotA plate 1**

- TotA Ta = 61.5ᴼC Rp49 Ta = 60ᴼC

- Standard curve efficiency = 106%

- Non-Rt and blank controls showed low/no amplification (range non-RT = 28-32CT, blank= 35-36CT)

- melt curves for TotA primer samples all good (77.17ᴼC) although Y,W samples in row A had slightly lower melting point (technical replicates in row E same as other samples)

**TotA plate 2**

- TotA Ta = 61.5ᴼC Rp49 Ta = 60ᴼC

- Standard curve efficiency = 96%

- Non-Rt and blank controls showed low/no amplification (range non-RT= 30-33CT, blank= 34-35CT)

- melt curves for TotA primer samples all good (76.43ᴼC) again Y,W samples in row A had slightly lower melting point

**UPD3 plate 1** (labelled plate 4 in folder as 4th upd3 plate run)

- UPD3 Ta = 60ᴼC Rp49 Ta= 60ᴼC

- Standard curve efficiency = 106%

- Non-Rt and blank controls showed low/no amplification (range non-RT = 32-37CT, blank= 33CT)

- melt curves for UPD3 primer samples all good (77.18ᴼC)

**UPD3 plate 2** (labelled plate 5 in folder as 5th upd3 plate run)

- UPD3 Ta = 60ᴼC Rp49 Ta= 60ᴼC

- Standard curve efficiency = 110%

- Non-Rt and blank controls showed low/no amplification (range non-RT = 35-36CT, blank= 36-37CT)

- melt curves for UPD3 primer samples all good (77.03ᴼC)

***Figure S3: Experimental design* (I)** Design of experiments to assay **(1)** survival (n= 15 vials of 15-17 flies/vial/infection treatment/sex/fly line) and **(2)** internal bacterial load following systemic bacterial infection with *Pseudomonas entomophila* to test the role of the Jak/Stat pathway (n= 15 vials of 8-10 flies/vial/infection treatment/sex/fly line) on **(3)** disease tolerance showing host vigor (good health state), more tolerant and less tolerant health state. Each point in disease tolerance (3) panel represents replicate-matched data [n= 15 vials/infection treatment/sex/fly line – with each vial containing 25 flies] from survival (1) and bacterial load (2). **(II)** A split-vial experimental design, where after infection each vial containing 25 flies of each treatment, sex and fly line combination were divided into 2 vials for measuring **(A)** survival following infection (15-17 flies/treatment) and **(B)**. internal bacterial load (8-10 flies/treatment).

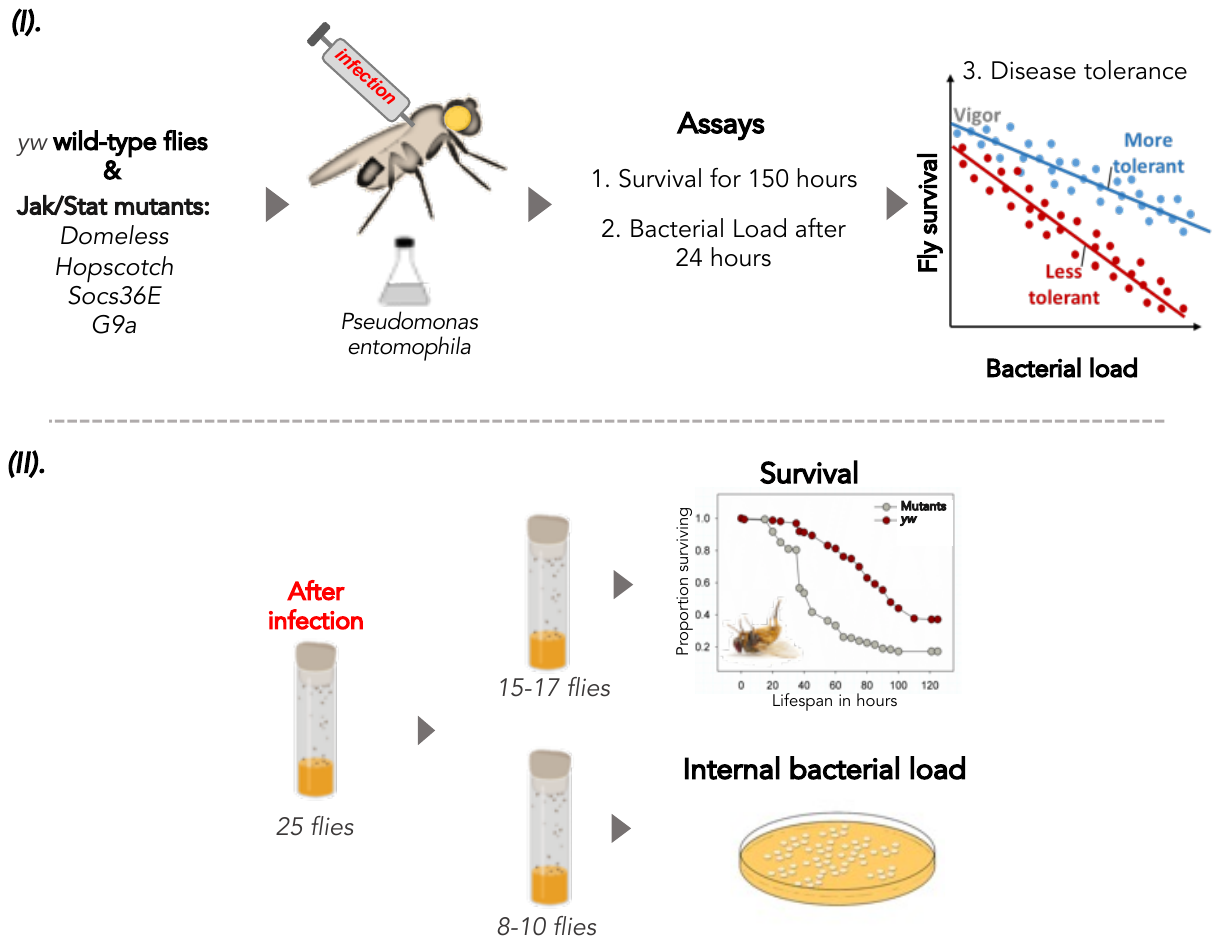

***Table S2*.** Summary of mixed effects Cox model, for wildtype *yw* and flies without Jak/Stat pathway components during systemic *P. entomophila* infection. We used data from the individuals of 3-day adult males and females infected with OD_600_ = 0.05 infection dose of *P. entomophila* for each fly lines and specified the model as: survival ~ treatment x sex x fly line (1|vial), with ‘treatment’, ‘sex’ and ‘fly line’ as fixed effects, and ‘vials’ as a random effect. The table shows model output (ANOVA) for survival post-infection for flies with fully function immune system and lacking Jak/Stat - pathway.

|  | ***Source*** | ***Log lik.*** | ***Chi sq.*** | ***df*** | ***p*** |
| --- | --- | --- | --- | --- | --- |
| ***Wildtype vs. outcross*** | Treatment (SI vs I) | -5956.3 | 500.18 | 1 | <0.001 |
|  | Sex | -5953.8 | 5.0844 | 1 | 0.38 |
|  | Fly line | -5945.8 | 15.929 | 1 | 0.54 |
|  | Treatment x sex | -5945.7 | 0.3030 | 1 | 0.58 |
|  | Treatment x fly line | -5945.7 | 0.0077 | 1 | 0.93 |
|  | Sex x fly line | -5945.3 | 0.7848 | 1 | 0.37 |
|  | Treatment x sex x fly line | -5942.8 | 4.9490 | 1 | 0.02 |
|  | *Random effect* | *Std dev* |  |  |  |
|  | *vial* | *0.61* |  |  |  |

***Table S3*.** Summary of mixed effects Cox model, for each fly line. We used data from the individuals of 3-day adult males and females infected with OD_600_ = 0.05 infection dose of *P. entomophila* for each fly line and specified the model as: survival ~ treatment x sex x fly line (1|vial), with ‘treatment’, ‘sex’ and ‘fly line’ as fixed effects, and ‘vials’ as a random effect. The table shows model output (ANOVA) for survival post-infection for flies with fully function immune system (*yw*) and flies lacking Jak/Stat – pathway components.

|  | ***Source*** | ***Log lik.*** | ***Chi sq.*** | ***df*** | ***p*** |
| --- | --- | --- | --- | --- | --- |
|  | Treatment | -30378 | 1042.2 | 1 | <0.001 |
|  | Sex | -30357 | 42.39 | 1 | <0.001 |
|  | Fly line | -29746 | 1222.9 | 4 | <0.001 |
|  | Treatment x sex | -29746 | 0.050 | 1 | 0.82 |
|  | Treatment x fly line | -29744 | 2.732 | 4 | 0.60 |
|  | Sex x flyline | -29736 | 15.57 | 4 | 0.003 |
|  | Treatment x fly line x sex | -29734 | 4.506 | 4 | 0.34 |
|  | *Random effect* | *Std dev* |  |  |  |
|  | *vial/block* | *0.49* |  |  |  |

***Table S4*.** Summary of non-parametric one-way ANOVA (Kruskal-Wallis test) of effects on bacterial load data measures 24 hours following OD_600_=0.05 *P. entomophila* systemic infection for wildtype (*yw)* flies and transgenic lines. We analysed the load data (log_10_ transformed) by fitting ‘fly line’ as categorical fixed effects for males and females separately.

| ***K-W*** | ***Sex*** | ***Source*** | ***DF*** | ***Chi Sq.*** | ***P*** |
| --- | --- | --- | --- | --- | --- |
|  | Female | Fly line | 4 | 9.1704 | 0.057 |
|  | Male | Fly line | 4 | 9.4565 | 0.0506 |

***Table S5.*** Summary of mixed effects Cox model, for *G9a* Jak/Stat pathway components during systemic *P. entomophila* infection. We used data from the individuals of 3-day adult males and females infected with OD_600_ = 0.05 infection dose of *P. entomophila* for each fly lines (*G9a*) and specified the model as: survival ~ treatment x sex x fly line (1|vial), with ‘treatment’, ‘sex’ and ‘fly line’ as fixed effects, and ‘vials’ as a random effect. The table shows model output (ANOVA) for survival post-infection for flies with fully function immune system and lacking *G9a*.

|  | ***Source*** | ***Log lik.*** | ***Chi sq.*** | ***df*** | ***p*** |
| --- | --- | --- | --- | --- | --- |
| ***G9a*** | Treatment | -8485.2 | 310.1 | 1 | <0.001 |
|  | Sex  Fly line | -8481.2  -8436.6 | 8.07  89.1 | 1  1 | 0.004  <0.001 |
|  | Treatment x sex  Treatment x fly line  Sex x fly line  Treatment x sex x fly line | -8436.5  -8433.9  -8426.3  -8426.3 | 0.22  5.13  15.2  0.00 | 1  1  1  1 | 0.63  0.002  <0.001  1.0 |
|  | *Random effect* | *Std dev* |  |  |  |
|  | *vial/block* | *0.25* |  |  |  |

***Table S6.*** Summary of non-parametric one-way ANOVA (Kruskal-Wallis test) of effects on bacterial load data measured at 24 hours following OD_600_=0.05 *P. entomophila* systemic infection for *G9a* transgenic lines. We analysed the load data (log_10_ transformed) by fitting ‘fly line’ (*G9a^+/+^ and G9a^-/-^*) as categorical fixed-effects for males and females separately.

| ***Fly line*** | ***Sex*** | ***S*** | ***Z*** | ***p*** |
| --- | --- | --- | --- | --- |
| ***G9a*** | Female | 641.5 | 3.065 | 0.002 |
|  | Male | 548.5 | 1.479 | 0.139 |
